# Characterisation of the *in-vivo* miRNA landscape in *Drosophila* ribonuclease mutants reveals Pacman mediated regulation of the highly conserved *let-7* cluster during apoptotic processes

**DOI:** 10.1101/2023.05.30.542828

**Authors:** Elisa I. M. Bernard, Benjamin P. Towler, Sarah F. Newbury

## Abstract

The control of gene expression is a fundamental process essential for correct development and to maintain homeostasis. Many post-transcriptional mechanisms exist to maintain the correct levels of each RNA transcript within the cell. Controlled and targeted cytoplasmic RNA degradation is one such mechanism with the 5’-3’ exoribonuclease Pacman (XRN1) and the 3’-5’ exoribonuclease Dis3L2 playing crucial roles. Loss of function mutations in either Pacman or Dis3L2 have been demonstrated to result in distinct phenotypes, and both have been implicated in human disease. One mechanism by which gene expression is controlled is through the function of miRNAs which have been shown to be crucial for the control of almost all cellular processes. Although the biogenesis and mechanisms of action of miRNAs have been comprehensively studied, the mechanisms regulating their own turnover are not well understood. Here we characterise the miRNA landscape in a natural developing tissue, the *Drosophila melanogaster* wing imaginal disc, and assess the importance of Pacman and Dis3L2 on the abundance of miRNAs. We reveal a complex landscape of miRNA expression and show that whilst a null mutation in *dis3L2* has a minimal effect on the miRNA expression profile, loss of Pacman has a profound effect with a third of all detected miRNAs demonstrating Pacman-sensitivity. We also reveal a role of Pacman in regulating the highly conserved *let-7* cluster (containing *miR-100, let-7* and *miR-125*) and present a genetic model outlining a positive feedback loop regulated by Pacman which begins to explain the apoptotic phenotype observed in Pacman mutants.

## INTRODUCTION

MicroRNAs (miRNAs) are a class of small non-coding RNAs that serve as regulators of gene expression in eukaryotic cells. miRNAs are 20-22 nucleotide (nt) long transcripts which have been shown to regulate the expression of a diverse range of RNAs and play a role in virtually every aspect of development. Since each miRNA can target many mRNAs, and mRNAs can include binding sites for a number of miRNAs, the impact of miRNA activity on the transcriptome and cellular function is profound and complex (Brennecke et al. 2003; Ge et al. 2012; Zhang and Cohen 2013; Aparicio et al. 2015; Bolin et al. 2016; Kane et al. 2018; Volin et al. 2018; Yu et al. 2020; Bejarano et al. 2021). In addition, dysregulation of a single miRNA can result in dramatic phenotypes in model organisms such as *Drosophila* and mice (Schertel et al. 2012; Towler et al. 2015; Bartel 2018). Furthermore, altered miRNA expression is known to contribute to the pathogenesis of human diseases including cancer (Mendell and Olson 2012).

miRNAs are usually transcribed as long primary transcripts (pri-miRNAs) which undergo a series of processing events to produce mature miRNAs. The genomic location of miRNAs shows great variety; they can be localised within genes encoding proteins, snoRNAs, lncRNAs, tRNAs or in separate transcription units and are often present in clusters containing multiple miRNAs of the same family (Soleimani et al. 2020). In the canonical pathway, pri-miRNAs form hairpin structures which are cleaved co-transcriptionally by the microprocessor complex comprising DGCR8 (Pasha in invertebrates) and Drosha to release the pre-miRNA hairpin loop which is then exported to the cytoplasm. Dicer (Dicer-1 in *Drosophila*) cleaves the hairpin to produce the miRNA duplex, which comprises two imperfectly complementary strands of around 22nt. Finally, one miRNA strand of the miRNA is loaded onto an Argonaute protein (Ago; Ago1 in *Drosophila*) which is part of the miRNA-induced silencing complex (RISC). Alternatively, miRNAs (termed mirtrons) can originate from introns of protein-coding genes (Ruby et al. 2007). After splicing of the mirtron precursor, the intron lariat is de-branched to allow folding and loading onto Dicer. Some mirtrons, termed “tailed mirtrons”, are derived from longer pri-miRNAs which need to be trimmed before Dicer binding and cleavage (Flynt et al. 2010). The processed mirtron is then loaded onto an Ago protein to provide an active complex, which then interacts with the target RNA.

miRNAs have been reported to be extremely stable and exhibit slower turnover that most RNAs. This is thought to be because the 5’ and 3’ ends of each miRNA are buried within the AGO protein, shielding them from ribonucleases (Sheu-Gruttadauria et al. 2019a; Sheu-Gruttadauria et al. 2019b). However, miRNAs with short half-lives have been observed and rates of turnover can also vary in different tissues and during transitions. For example, metabolic labelling of *Drosophila* S2 cells has shown that half-lives can vary from < 2 hours (e.g. *miR-12-5p*) to more than 24 hours (e.g. *bantam-3p*) (Reichholf et al. 2019). These data suggest that there are specific mechanisms which promote miRNA degradation in response to cellular or developmental cues.

While much is known about the regulation of miRNA biogenesis and activity, miRNA degradation pathways are comparatively poorly understood. The best understood pathway is target-directed miRNA degradation (TDMD) where another RNA (such as the lncRNA *CYRANO*) forms extensive base-pairing to miRNA, resulting in its degradation (Han et al. 2020; Shi et al. 2020). Recent studies have shown that the ZSWIM8 ubiquitin ligase (Dora in *Drosophila*) recognises Ago:miRNA complexes engaged with TDMD targets resulting in ubiquitination of Ago and degradation by the proteasome (Han et al. 2020; Shi et al. 2020). The miRNA is then degraded by unknown ribonucleases. If 3’ends of the miRNAs become exposed, such as by TDMD engagement or mutation of the PAZ domain, the miRNAs can become tailed by terminal nucleotidyl transferases such as TUT4 and TUT7 and subsequently trimmed by 3’-5’ exoribonucleases such as DIS3L2 (Heo et al. 2009; Ameres et al. 2010; Cazalla and Steitz 2010; Heo et al. 2012; Chang et al. 2013; Ustianenko et al. 2013; Sheu-Gruttadauria et al. 2019a). However, this tailing does not necessarily lead to miRNA degradation (Yang et al. 2020). An example of tail-mediated miRNA decay includes maternally deposited miRNAs in *Drosophila*, which can be cleared during the maternal to zygotic transition in a process that involves adenylation by the non-canonical polyA polymerase Wispy (Lee et al. 2014). Tailing and trimming of miRNAs has also been implicated in regulating their turnover by DIS3L or DIS3L2 which was shown to have significant functional importance in human cells (Shukla et al. 2019).

Despite their importance in post-transcriptional regulation, little is known about the exoribonucleases responsible for degrading specific miRNAs. Previous studies in *C. elegans* have shown that the nuclear exoribonuclease XRN2 facilitates miRNA release from RISC and degradation in a 5’-3’ direction (Chatterjee and Grosshans 2009). Cytoplasmic Xrn-1 was also shown to be involved in mature miRNA turnover in *C. elegans* irrespective of whether the miRNA was associated with Ago (Chatterjee et al. 2011). It is possible that XRN-1 is recruited to the miRNAs by the decapping scavenger enzyme DCS-1 (Bosse et al. 2013). In *Drosophila* wing imaginal discs, specific miRNAs are sensitive to depletion of the 5’-3’ exoribonuclease XRN1 (Pacman) or the 3’-5’ exoribonuclease Dis3 (*miR-277-3p* (Jones et al. 2013) and *miR-252-5p* (Towler et al. 2015) respectively). Therefore, further work is required to improve understanding of the specific roles ribonucleases play in the degradation of miRNAs.

Here, we have investigated whether miRNAs are sensitive to degradation by the 5’-3’ exoribonuclease Xrn1 (Pacman) or the 3’-5’ exoribonuclease Dis3L2. We characterise the *in vivo* miRNA landscape using *Drosophila* wing imaginal discs and assess their susceptibility to Pacman or Dis3L2 null mutations. We show that nearly a third of detected miRNAs are sensitive to Pacman expression and demonstrate a potential role for Pacman in regulating the highly conserved and developmentally important *let-7* miRNA cluster.

## RESULTS

### Global expression analysis characterises the microRNA landscape in *Drosophila* wing imaginal discs

Our previous work sought to explore the specific roles of ribonucleases in regulating development and cellular homeostasis. Using the *Drosophila melanogaster* imaginal disc we have shown that disruption of the major cytoplasmic decay complexes results in contrasting phenotypes (Jones et al. 2013; Waldron et al. 2015; Jones et al. 2016; Towler et al. 2016; Towler et al. 2020). Depletion of the 5’-3’ exoribonuclease Pacman (XRN1) results in increased developmental delay and apoptosis (Waldron et al. 2015; Jones et al. 2016) whereas depletion of Dis3L2 results in enhanced proliferation (Towler et al. 2020). Using RNA-sequencing we have identifed some of the ribonuclease sensitive transcripts that underpin the mutant phenotypes observed. *dilp8* is upregulated in *pacman* mutants and drives developmental delay (Waldron et al. 2015; Jones et al. 2016) whereas *idgf2* contributes to tissue overgrowth in *dis3L2* mutants (Towler et al. 2020). To provide a comprehensive assessment of the role of Pacman and Dis3L2 in regulating miRNAs, which also may contribute to the observed phenotypes, we performed small RNA sequencing on Pacman null (*pcm^14^*) and Dis3L2 null (*dis3L2^12^*) mutant wing imaginal discs from wandering L3 larvae along with their isogenic control lines (*pcm^WT^* and *dis3L2*^WT^)(Waldron et al. 2015; Towler et al. 2020).

We first used this data as a starting point to reveal, for the first time, the overall miRNA expression landscape, in terms of numbers, expression levels and genomic origin of miRNAs in wild-type *Drosophila* wing imaginal discs. We identify 201 miRNAs out of 481 annotated *Drosophila* miRNAs (miRbase release 6 v22, (Kozomara et al. 2018)) as confidently and consistently expressed in these WIDs across all 8 control replicates). Expression of miRNAs in these controls were highly correlated (R^2^= 0.997; p<0.0001; Supplemental Fig S1A-B) demonstrating the technical reliability and consistency of miRNA expression within these tissues. First, we assessed the genomic origin of the detected miRNAs and observed 34% of these originated from exons within host transcripts (Fig 1A). Only 17% were expressed from their own locus whilst 32% and 16% of miRNAs respectively were derived from large introns or were mirtrons (Fig 1A). Consistent with a role of lncRNAs as miRNA host transcripts, 27% of miRNAs are processed from lncRNAs and 96% of these are encoded within exons (Fig 1B-D). Strikingly, 54% of miRNAs were hosted within mRNAs, with the majority of these within intronic regions (86%) and all but 1 of those that derived from mRNA exons were located in the 3’ UTR. This outlines the extensive regulation required to maintain miRNA homeostasis.

**Figure 1:**
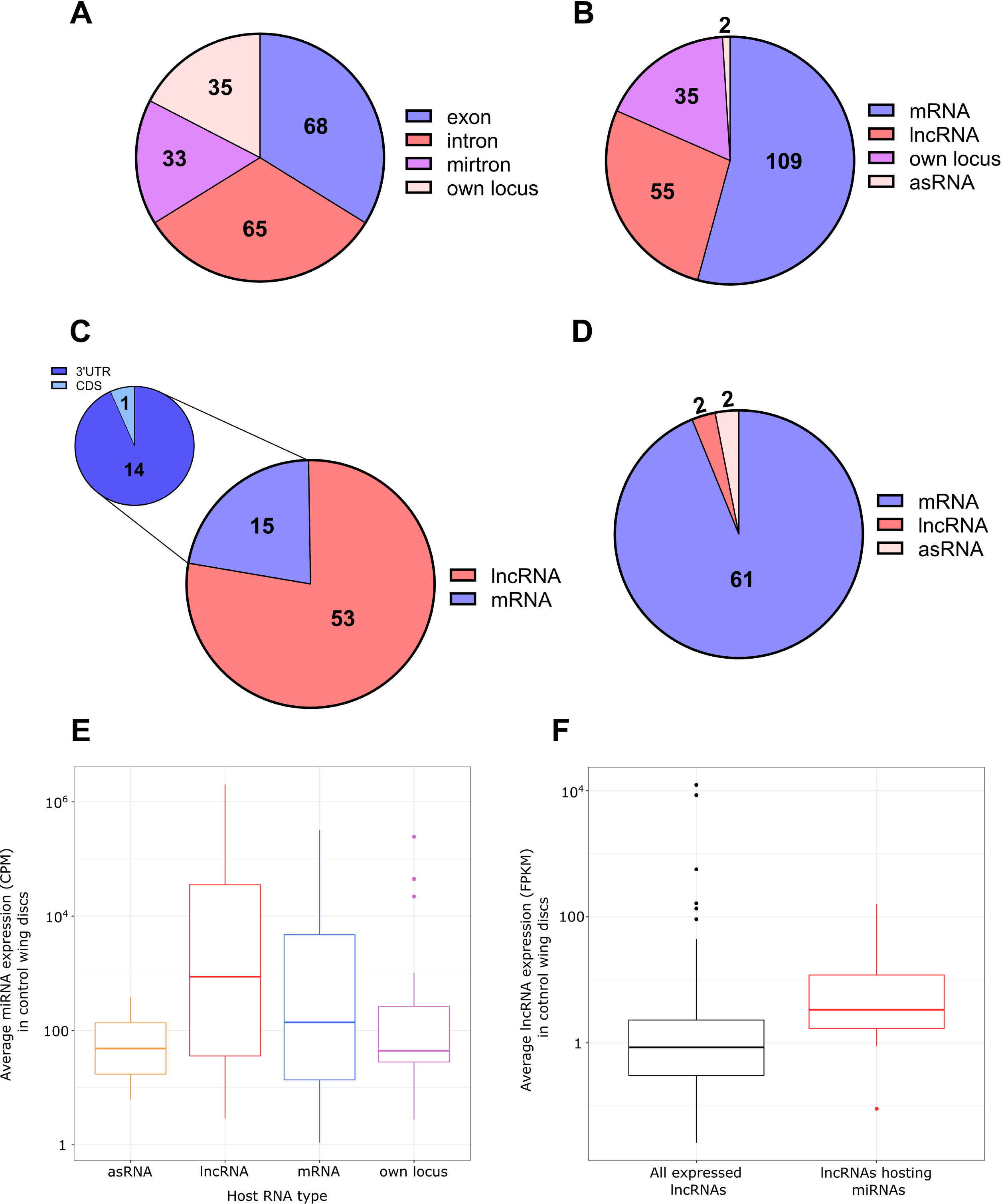
Overview of the miRNAs expressed in wing imaginal discs. **A)** Genomic origin of all miRNAs reliably detected in all control replicates. **B)** Proportion of reliably detected mRNAs by the nature of their host RNA. **C)** miRNAs derived from exons within a host transcript tend to derive from lncRNAs. Those from mRNAs tend to reside in the 3’UTR of their host. **D)** Most miRNAs derived from intronic regions within a host RNA are encoded within mRNAs. **E)** miRNAs derived from host RNAs tend to be more abundant than those transcribed from their own locus. **F)** lncRNAs encoding miRNAs are more abundant than the average expression of lncRNAs in the wing disc. 508 lncRNAs in total detected in wing imaginal discs from Towler *et al*. p=0.001253.

Next we asked if miRNAs from different host transcripts or genomic origins demonstrated expression patterns. We saw that, on average, those derived from host transcripts were more abundant that those from their own locus (Fig 1E, Supplemental Fig S1C) and interestingly, lncRNAs hosting miRNAs were both more abundant and longer than the genome average as calculated from our previous data (Towler et al. 2020) (Fig 1F, Supplemental Fig S1D).

### Loss of Dis3L2 has a minimal effect on the miRNA landscape

Having characterised the miRNA transcriptome in control tissues we then aimed to assess the effect of Dis3L2 loss of function on miRNA expression. Previous work has used mammalian cell lines to demonstrate a role for DIS3L2 in miRNA and pre-miRNAs decay following their terminal tailing including uridylation by uridyltransferases such as TUT4 and TUT7 (Heo et al. 2009; Heo et al. 2012; Chang et al. 2013; Ustianenko et al. 2016; Shukla et al. 2019). However, a genome-wide study on the effect of Dis3L2 on mature miRNAs in a natural tissue has not been performed. We compared the miRNA expression profile in *dis3L2^WT^* to that in *dis3L2^12^* null mutant wing imaginal using two analysis pipelines, miRDeep2 (Friedlander et al. 2012) and sRNAbench (Aparicio-Puerta et al. 2019; Aparicio-Puerta et al. 2022) with a miRNA requiring a significant change in expression by both pipelines to be considered ribonuclease-sensitive (>1.5 fold change, padj<0.05). A total of 221 miRNAs were detected in all *dis3L2^WT^* and *dis3L2^12^* replicates. All processed data can be found in Supplemental File S1.

Loss of *dis3L2* resulted in minimal changes in the miRNA expression profile with only 13 (6%) showing upregulation and 10 (5%) showing downregulation in *dis3L2^12^* tissues (Fig 2A-C). Whilst we observed an increase in expression of miRNAs hosted within asRNAs, this is likely biased by the small number of these miRNAs expressed (2 in total). Therefore, we did not observe any strong trend in miRNA sensitivity to Dis3L2 based on their origin or host RNA (Supplemental Fig S2A, C-D). As we saw an absence of miRNAs transcribed from their own locus in the downregulated pool, we used our previous data set to assess correlation between the sensitivity of the miRNA or its host RNA to Dis3L2; all but 1 miRNA changed in levels independently from their host suggesting these miRNA changes represent independent post-transcriptional regulation rather than a result of regulation of their host transcript (Supplemental Fig S2E).

**Figure 2:**
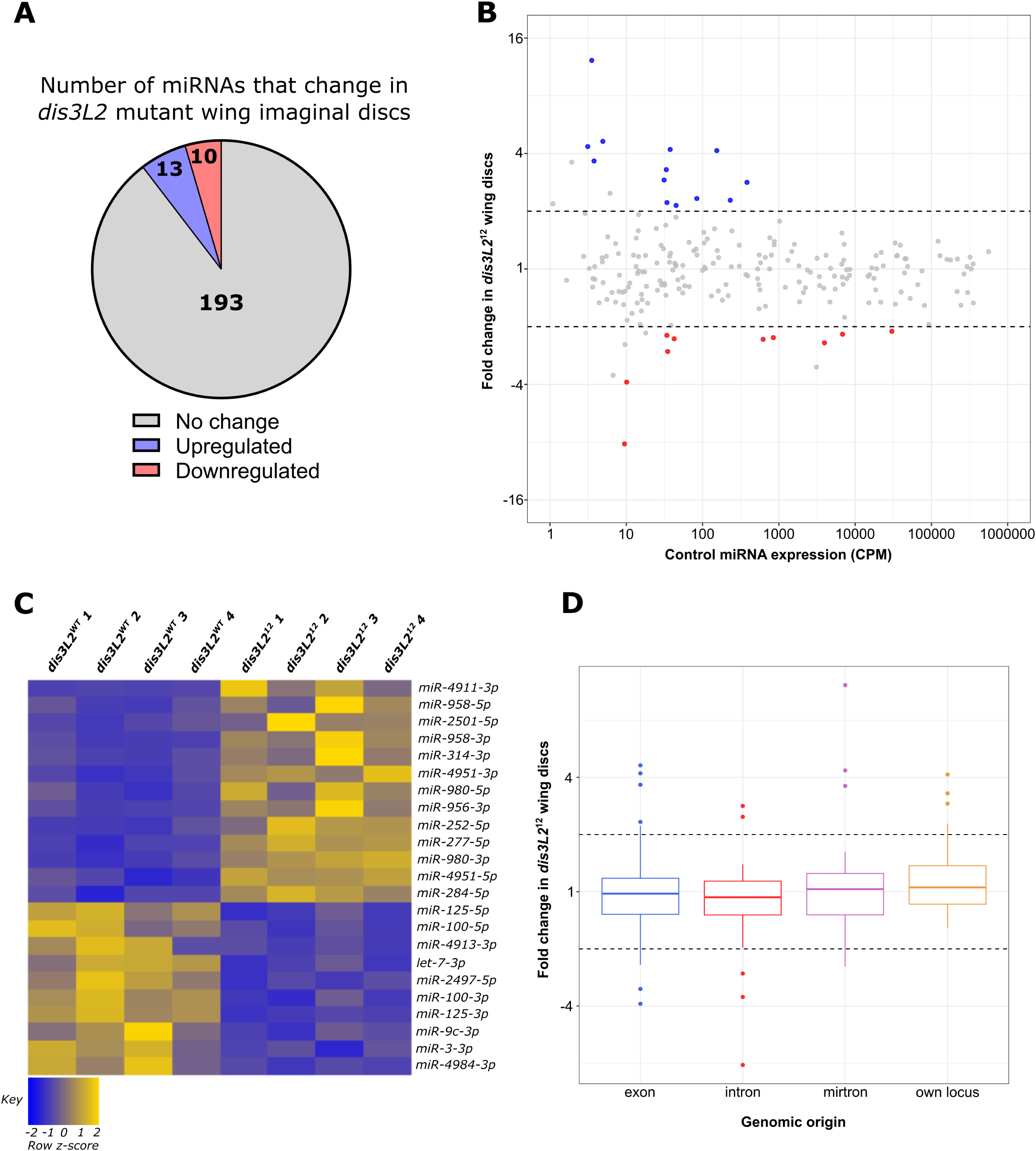
A specific subset of miRNAs are affected by the loss of Dis3L2. **A)** Proportion of miRNAs significantly dysregulated in *dis3L2^12^*WIDs. miRNAs determined as differentially expressed if the fold change is >2 and called significant by both analysis pipelines. **B)** MA plot displaying all miRNAs expressed WIDs with upregulated miRNAs in blue and downregulated in red. Dotted lines represent ^+^2 fold change. Displayed CPM and fold change calculated using DESeq2 **. C)** Heatmap representing the expression of differentially expressed miRNAs in all replicates of *dis3L2^WT^*and *dis3L2^12^* wing imaginal discs. **D)** Boxplot displaying fold change of all miRNAs in *dis3L2^12^* wing discs by genomic origin. Dotted lines represent ^+^ 2 fold change.

### Pacman function is required to maintain the miRNA expression profile

Our previous work, together with that of others in *C. elegans* and human cells, have suggested an important role of Pacman in miRNA regulation (Bail et al. 2010; Chatterjee et al. 2011; Jones et al. 2013). To determine the extent that Pacman regulates the miRNA landscape, and to gain insights into the ways in which these miRNAs may induce the phenotypes observed, we used the above miRNA-seq experiment to compare the levels of miRNAs between *pcm^14^*mutants and the isogenic controls. Similarly to our analysis with *dis3L2* mutant tissues we used two independent pipelines to assess differential expression between *pcm^WT^* and *pcm^14^* wing imaginal discs. Strikingly, we observed altered expression in 32% (70) of expressed miRNAs, with 80% of these demonstrating stabilisation in *pcm^14^* null tissues (Fig 3A-C). Interestingly, in contrast to *dis3L2* mutants, we observed a global trend of miRNAs transcribed from their own genomic locus displaying sensitivity to Pacman resulting in an enrichment of these miRNAs in the upregulated pool (Fig 3D, Supplemental Fig S2B-D). As with our analysis of Dis3L2-sensitive miRNAs, those miRNAs displaying Pacman sensitivity which were derived from host transcripts, largely showed regulation independent to that of their host (Supplemental Fig S2F).

**Figure 3:**
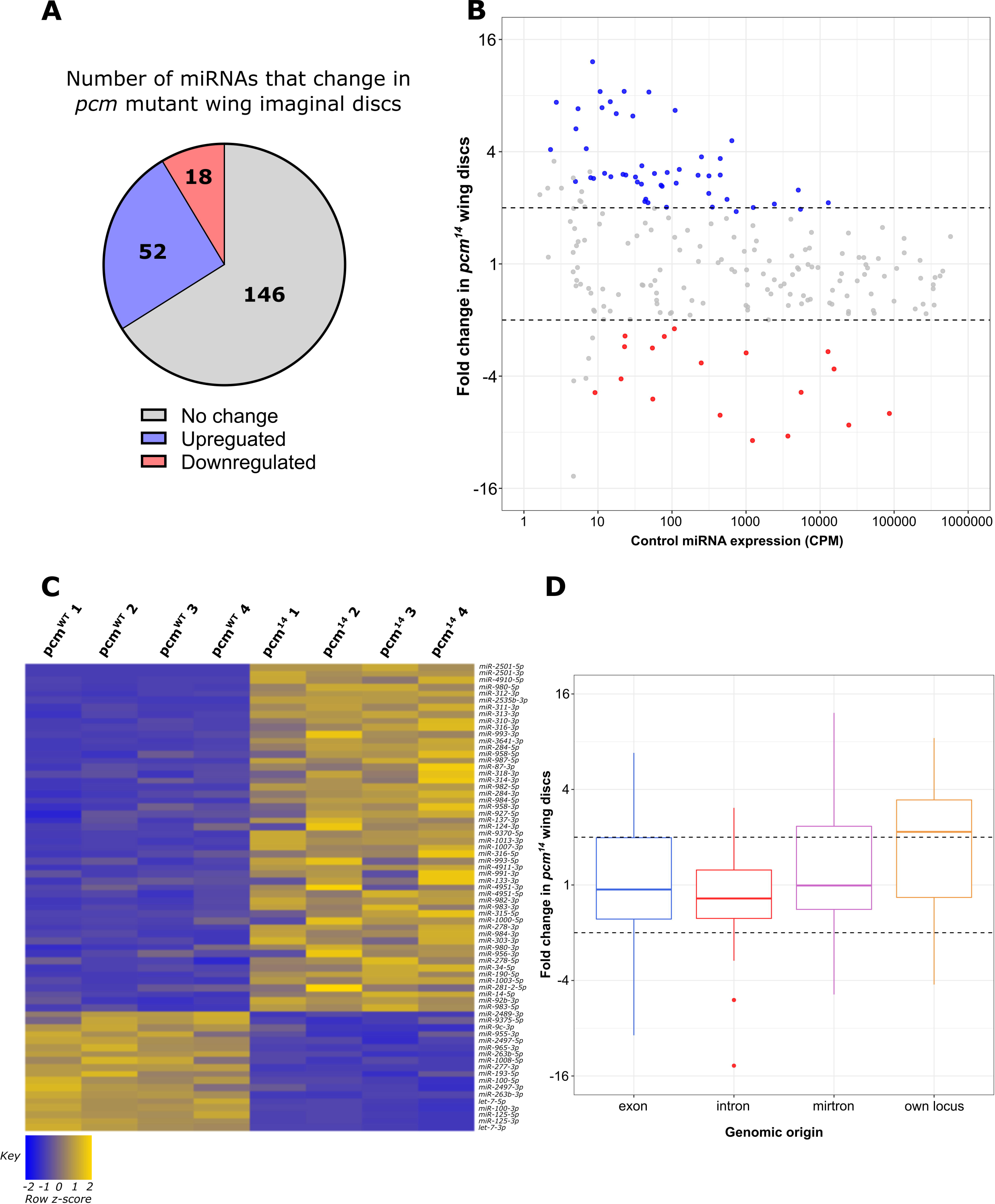
Loss of pcm results in changes in the miRNA landscape. **A)** 30.8% of all miRNAs display significantly differential expression in *pcm^14^* WIDs. miRNAs determined as differentially epxressed if fold change >2 and called significant by both analysis pipelines**. B)** MA plot displaying all miRNAs expressed WIDs with upregulated miRNAs in blue and downregulated in red. Dotted lines represent ^+^2 fold change. Displayed CPM and fold change calculated using DESeq2. **C)** Heatmap representing the expression of differentially expressed miRNAs in all replicates of *pcm^WT^* and *pcm^14^* wing imaginal discs. **D)** Boxplot displaying fold change of all miRNAs in *pcm^14^* wing discs by genomic origin. Dotted lines represent ^+^2 fold change.

### Pacman sensitive miRNAs do not appear to be in Ago complexes

Having observed a strong impact of Pacman mutations on the miRNA landscape we decided to determine whether selected upregulated miRNAs were likely to be functional (i.e Ago bound within RISC), or if they were in a free miRNA pool awaiting degradation. First, we sought to validate our miRNA-sequencing using TaqMan qRT-PCR using newly dissected WIDs. Here, we selected miRNAs displaying differential expression in *pcm* or *dis3L2* mutant tissues. Whilst only *miR-4911-3p* was shown to be differentially expressed in *dis3L2^12^* tissues (Fig 4A), all selected miRNAs demonstrating Pacman-sensitivity were successful validated suggesting confidence in this dataset (Fig 4B). *miR-2501-5p* was the most upregulated miRNA in *pcm^14^* tissues, and interestingly has previously been shown to be a tailed mirtron (Flynt et al. 2010) processed from an inton in its host gene *nulp1*. Therefore, we assessed the levels of its *pre-miRNA* in *pcm^14^* tissues which showed no change in expression (Fig 4B), thus suggesting the increase in *miR-2501-5p* was driven by defective post-transcriptional regulation in the absence of Pacman.

**Figure 4:**
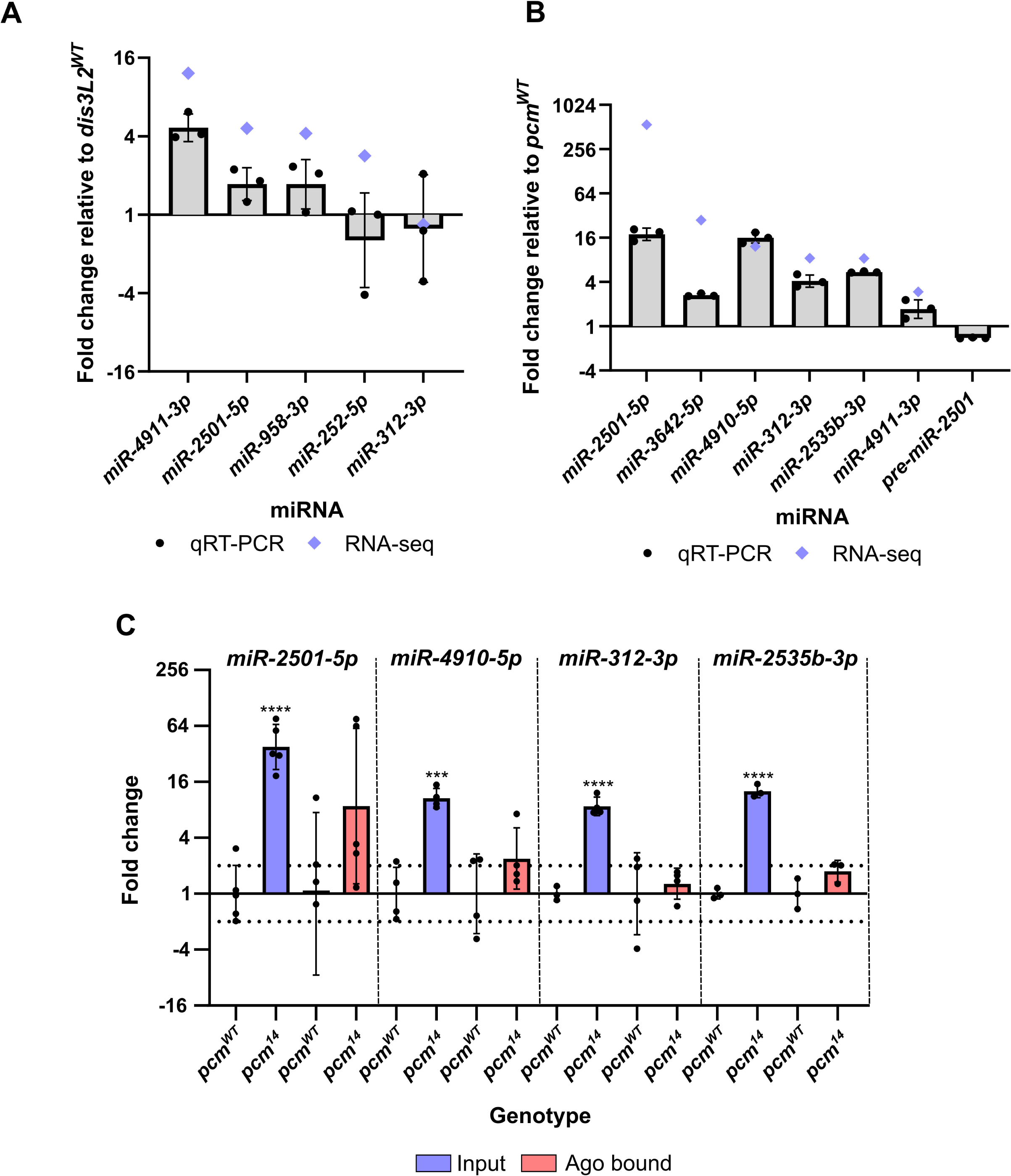
miRNAs differentially expressed in pcm^14^ wing imaginal discs are not RISC-associated. **A)** qRT-PCR validation of differentially expressed miRNAs in *dis3L2^12^* mutant tissues. Blue diamonds represent fold change by miRNA-seq. n=3, error bars represent SD, statistical information can be found in Table S1. **B)** qRT-PCR validation of differentially expressed miRNAs in *pcm^14^* mutant tissues. Blue diamonds represent fold change by miRNA-seq. n=3, error bars represent SD, statistical information can be found in Table S2. **C)** Assessment of Ago/RISC associated miRNAs. RNA from wing imaginal discs was isolated using the TraPR kit (Lexogen) and the differential expression between control and *pcm^14^* was assessed for input and Ago bound fractions. Differential expression was only identified in the input fraction suggesting the excess miRNA is not RISC associated. n≥3, error bars represent SD, ***p<0.001, ****p<0.0001, no stars=p>0.05. Full statistical information can be found in Table S3.

To determine if the selected miRNAs were functionally associated with an Ago protein we isolated Ago-miRNA complexes from *pcm^WT^* and *pcm^14^* wing imaginal discs using the TraPR kit from Lexogen. This allows the identification of small RNAs bound by Ago (Ago1 or Ago2) using ion exchange chromatography. Free RNA stays bound on the column while small RNAs protected by Ago flow through and therefore Ago-associated miRNAs can subsequently be measured by TaqMan qPCR. We validated the specificity of the kit for miRNAs by comparing the binding of a snoRNA, *snoR442*, with the selected miRNAs. Whilst all selected miRNAs were reliably detected by TaqMan qPCR, *snoR442* was not. We next compared the expression of our selected Pacman-sensitive miRNAs in the input and Ago-bound fractions and whilst the input fraction displayed the expected change in expression in *pcm^14^* tissues, we did not observe a significant increase in the amount of miRNA associated with Ago complexes (Fig 4C). This suggests that Pacman may degrade miRNAs that are no longer required for targeting and have been released from the Ago complex to await degradation.

### The developmentally important and conserved *let-7* cluster is Pacman-sensitive

Our analysis of the effect of Pacman depletion on the *in vivo* miRNA landscape has revealed that 80% of miRNAs which show differential expression in *pcm^14^* mutants are increased in expression, as would be expected when a ribonuclease is depleted. However, we also observed 18 miRNAs that displayed reduced expression in the absence of Pacman. Strikingly, 6 of these are located in the single, conserved *lncRNA:let7C.* This single lncRNA is the host of *miR-100, let-7* and *miR-125* (Sokol et al. 2008; Emmrich et al. 2014) with their downregulation in *pcm^14^* mutants ranging from −4.9-fold (*miR-100-5p*) to −8.9-fold (*let-7-3p*). The *let7C* lncRNA miRNA cluster is conserved between flies and humans with the relative order of miRNAs also conserved (Sokol et al. 2008; Emmrich et al. 2014). In *Drosophila*, these miRNAs are located within the single exon of the host lncRNAs, whilst in humans they are located in intron 3 (Fig 5A).

**Figure 5:**
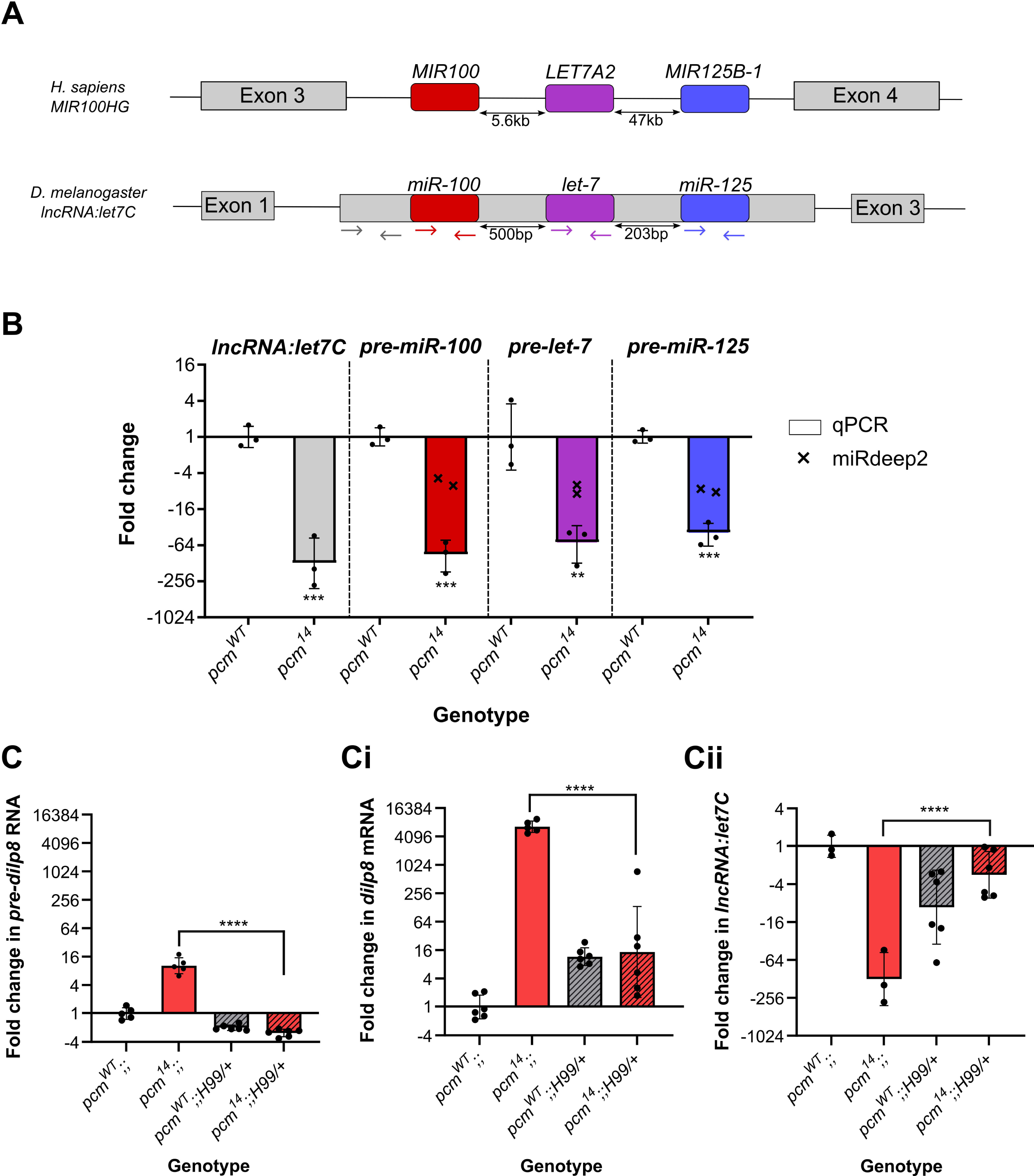
Increased apoptosis in pcm^14^ wing discs drives the changes in expression of the let7C cluster and dilp8. **A)** Genomic organisation of *let7C* demonstrating its conservation between humans and *Drosophila melanogaster*. Coloured arrows represent the location of the primers used in B. **B)** The *let7C* host lncRNA and pre-miRNAs within the host are downregulated in *pcm^14^* wing discs. miRdeep2 data (crosses) display the fold change in both arms (5p and 3p) of the mature miRNAs by miRNA-seq. **p<0.01, ***p<0.001, n=3, error bars represent SD. **C)** The differential expression of *pre-dilp8* (i) *dilp8* (ii) and *lncRNA:let7C* (iii) in *pcm^14^* wing discs is largely driven by apoptosis as inhibition of apoptosis (using *H99*) in *pcm^14^*tissues rescues the altered expression. ****p<0.0001, n≥3, error bars represent SD. Statistical information for all comparisons is shown in Table S4.

In *Drosophila*, null mutations in *let7C* result in lethality with a 43% penetrance, with surviving adults having defects in flight, mobility and fertility (Sokol et al. 2008). Studies on the phenotypic effects of the individual miRNAs within this complex showed that *let-7* is critical for the timing of cell cycle exit in wing imaginal discs (Reinhart et al. 2000; Moss 2007) as well as other phenotypes such as severe defects in female fertility and oviposition (Sokol et al. 2008). We therefore decided to further investigate the effect of Pacman depletion on *let7C* biology. First, we validated the change in each pre-miRNA hairpin by qRT-PCR and showed a strong downregulation (Fig 5B). This was expected as both arms (5p- and 3p-) of the miRNA are downregulated in our miRNA-seq data. Next, we asked if this was a result of downregulation of the host transcript, *lncRNA:let7C*, or if it was a dysregulation of pre-miRNA processing, again using qRT-PCR. Using primers specific for the lncRNA (Fig 5A) we demonstrated a strong reduction in *lncRNA:let7C*, consistent with that of the pre-miRNAs suggesting that this was indeed driven by a downregulation of the host lncRNA, likely driven by a reduction in transcription (Fig 5B).

One of the most studied targets of *let-7* and *miR-125* is *chronologically inappropriate morphogenesis* (*chinmo*)(Wu et al. 2012; Chawla et al. 2016; Pandey et al. 2021) which acts in a dosage-dependent manner to regulate the temporal identity and viability of neurones (Zhu et al. 2006; Chawla et al. 2016). To ask if the downregulation of *let-7* and *miR-125* is functional, we assessed *chinmo* expression in *pcm* mutant tissues and observed a significant increase in our previous RNA-seq data (Jones et al. 2016), which we validated as a 23.1-fold increase by qRT-PCR (Supplemental Fig S3). This therefore suggests that the reduction in *chinmo* required for a timely developmental transition is absent in *pacman* mutants as a result of decreased *let-7* and *miR-125* expression.

To further probe this pathway, we turned to our previous work, in which we showed that the insulin-like peptide Dilp8, which co-ordinates tissue growth with developmental timing, is upregulated in *pcm* mutants at the transcriptional and post-transcriptional levels (Jones et al. 2016). Dilp8 has previously been reported to be secreted from the imaginal discs in vesicle-like structures in response to tissue damage (Colombani et al. 2012; Garelli et al. 2012). This hormone then travels through the haemolymph to remote tissues such as the brain complex to suppress ecdysone production (Garelli et al. 2012; Colombani et al. 2015; Garelli et al. 2015). Lower levels of ecdysone production delay development and the onset of pupariation (Garelli et al. 2012; Colombani et al. 2015; Garelli et al. 2015). In *Drosophila* Kc-167 and S2 cultured cells, dissected organs and pre-pupae, the steroid hormone ecdysone (20-hydroxyecdysone) is required for expression of *let-7* and *let7C* (Sempere et al. 2002; Sempere et al. 2003; Chawla and Sokol 2012) although other regulatory factors may be involved (Bashirullah et al. 2003; Luhur et al. 2014).

Therefore, an attractive hypothesis is that the *pcm^14^* mutation results in upregulation of Dilp8 (as previously shown; (Jones et al. 2016)), resulting in lower levels of ecdysone in the hemolymph and reduced expression of *let7C* and its encoded miRNAs *miR-100*, *miR-125* and *let-7*. Since the involvement of Dilp8 in *let7C* regulation has not yet been tested, we decided to probe the potential relationship. We confirmed the systemic increase in *dlip8* and reduction in *lncRNA-let7C* expression throughout *pcm^14^*larvae (Fig 6A), and subsequently used the *GAL4-UAS* system to ectopically express wild-type *dilp8* throughout the wing disc (*69B-GAL4*). Using TaqMan qPCR, we demonstrated successful ectopic expression of *dilp8* RNA (Fig 6Bi), however, this in fact seemed to result in a slight increase in *let7C* expression rather than the reduction observed in *pcm^14^* tissues (Fig 6Bii). Therefore, the increase in *dilp8* expression alone is not sufficient to drive the observed reduction in *let7C*.

**Figure 6:**
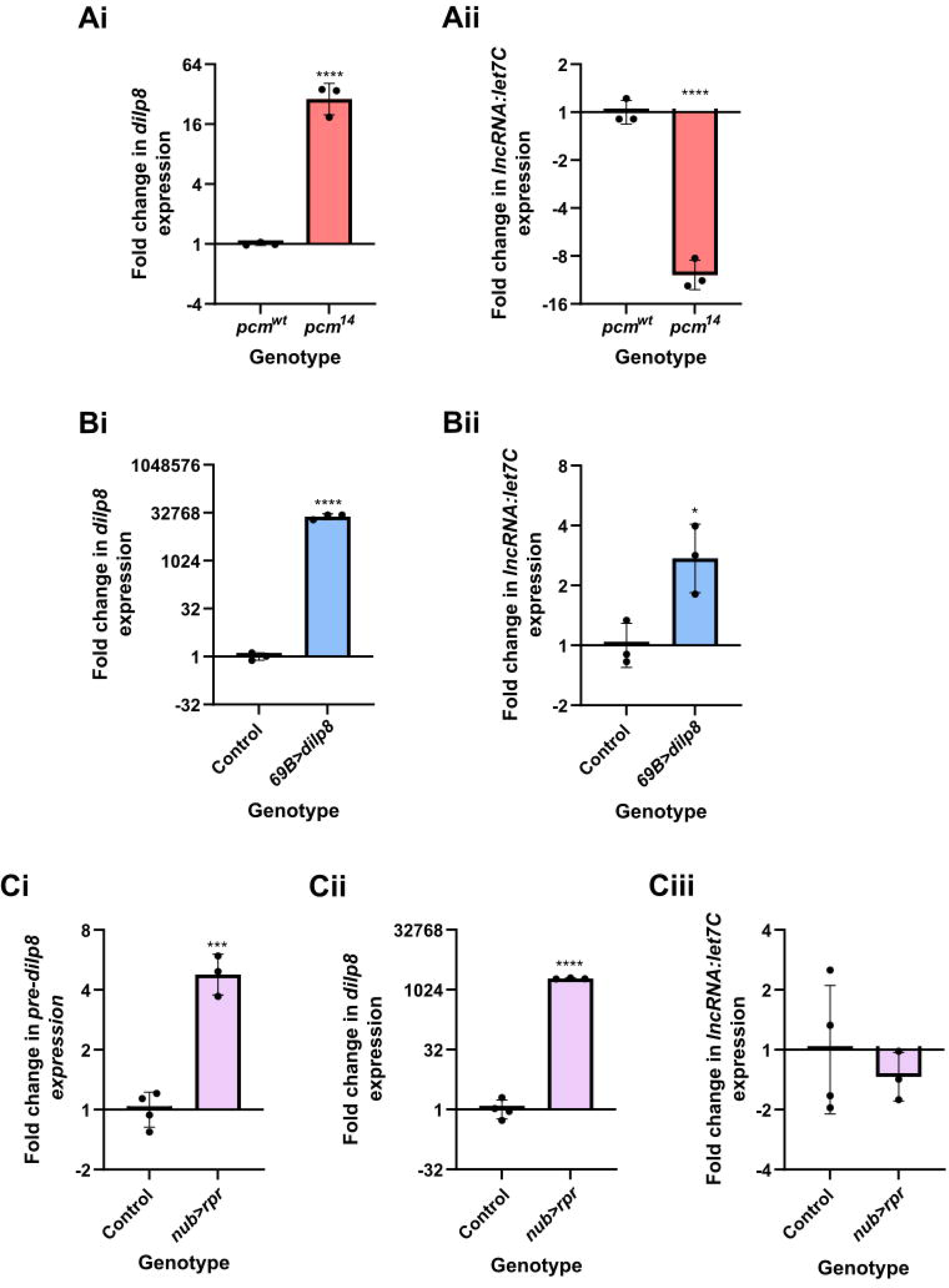
The induction of apoptosis or increased dilp8 expression alone is insufficient to drive downregulation of let7C. **A)** Increased expression of *dilp8* (i) and *let7C* (ii) can be observed throughout the L3 larvae. n=3, error bars represent SD, ****p≤0.0001. **B)** Overexpression of *dilp8* throughout the wing imaginal disc using *69B-GAL4* is not sufficient to drive *let7C* downregulation. (i) demonstrates *dlip8* expression and (ii) demonstrates *let7C* expression in wing imaginal discs. n=3, error bars represent SD, ****p<0.0001, *p=<0.05. **C)** Driving *UAS-rpr* in the wing pouch of the wing imaginal disc with *nub-GAL4* to induce apoptosis is sufficient to cause increased *pre-dilp8* (i) and *dilp8* (ii) expression but not the downregulation of *let7C*. n≥3, error bars represent SEM, ***p<0.001, ****p<0.0001. For all statistical information see Table S5.

Given that our previous work demonstrated a contribution of both transcriptional and post-transcriptional regulation to the increased *dlip8* expression in *pcm^14^* tissues (Jones et al. 2016), and that *dilp8* is secreted in response to tissue damage (Colombani et al. 2012; Garelli et al. 2012), we hypothesised that the apoptosis induced by loss of Pacman (Waldron et al. 2015) may contribute to the observed changes in *dlip8* and *let7C.* To address this we inhibited apoptosis using a heterozygous *H99* deletion which removes a single copy of the pro-apoptotic genes *hid, grim* and *rpr* (Waldron et al. 2015). When used as a heterozygote in *pcm^14^*mutants the *H99* deletion rescues the apoptotic phenotype (Waldron et al. 2015). Strikingly, the same genetic approach resulted in a complete rescue of *pre-dilp8*, *dlip8* and, crucially *let7C* (Fig 5C).

To determine whether apoptosis alone is sufficient for the observed changes in *dilp8* and *let7C* expression, or if specifically Pacman-induced apoptosis is required, we used *nubbin-GAL4* to ectopically express the pro-apoptotic *rpr* specifically in the wing pouch of the wing imaginal disc. This resulted in an increase in *pre-dilp8* and a substantial increase in mature *dlip8* expression, although to a lesser extent compared to in *pcm^14^* tissues, suggesting loss of Pacman may accentuate these effects. However, despite the increase in *pre-dilp8* and *dilp8* we did not observe a reduction in *let7C* expression (Fig 6C). Therefore, these data suggest that whilst apoptosis alone is sufficient to drive changes in *dilp8* expression, the loss of Pacman *and* apoptosis are responsible for the reduction in *let7C* expression demonstrating a crucial role for Pacman in regulating these developmentally important miRNAs.

Confirmation that Pacman regulates a complex network of proteins and RNAs which normally prevent apoptosis is supported by further inspection of our miRNA-seq data. As can be seen in Fig 3C and Supplemental File S1, *miR-277*, *miR-263a* and *miR-263b* are also down-regulated in *pcm^14^* wing imaginal discs. It has previously been shown that a decrease in *miR-125, 277* and *263a* and *b* results in increased *hid* expression because *hid* includes binding sites for these RNAs in its 3’UTR (Hilgers et al. 2010; Bejarano et al. 2021). In addition, the pro-apoptotic RNAs *reaper*, *hid* and *grim* increased in levels during apoptosis with *reaper* and *hid* RNAs increasing at the post-transcriptional level in *pcm* null mutants (Waldron et al. 2015; Jones et al. 2016). These results are entirely consistent with a positive feedback pathway that occurs in *pacman* mutants to increase apoptosis (Fig 7).

**Figure 7:**
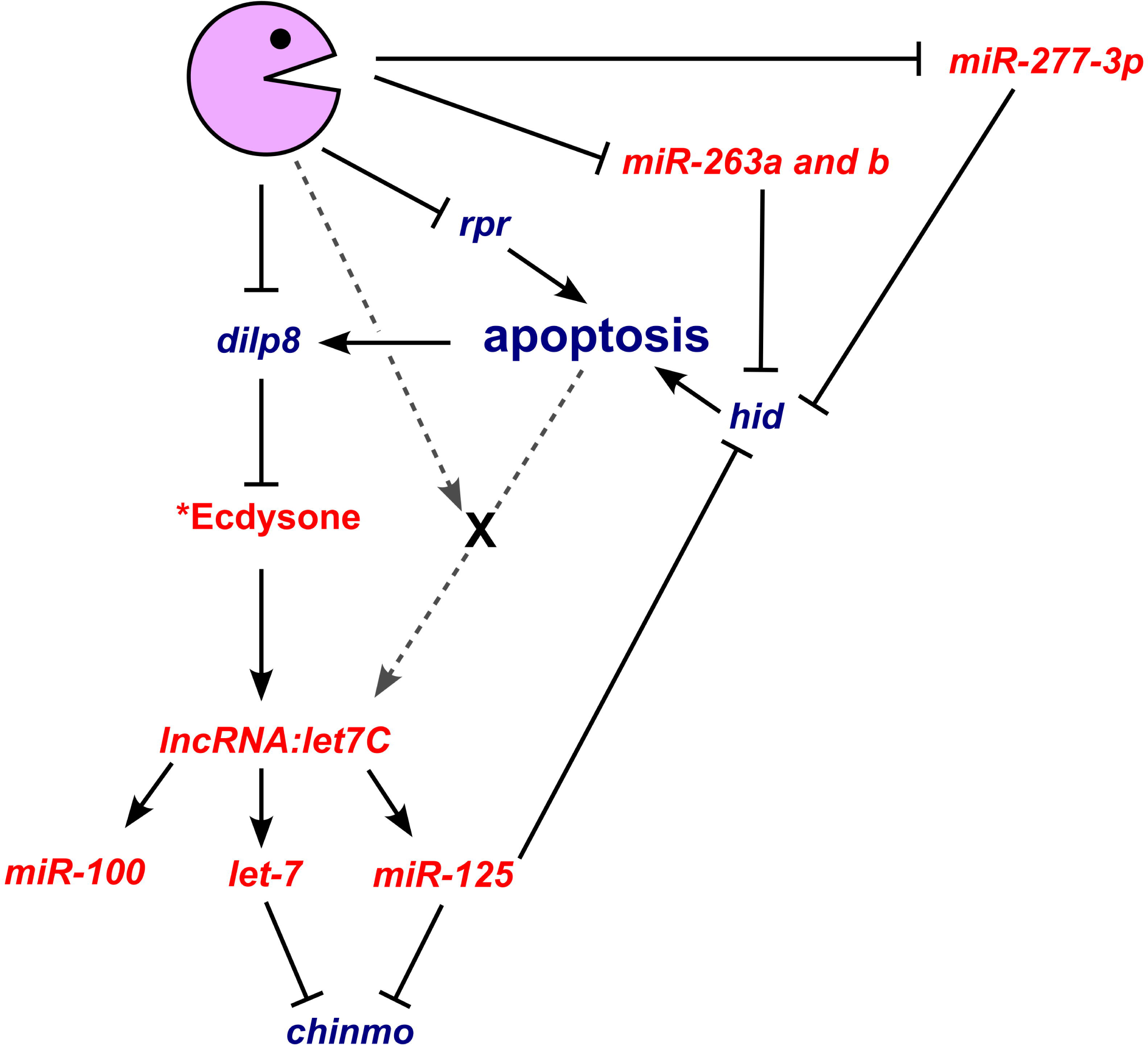
Genetic model whereby loss of Pacman results in the observed changes in gene expression and subsequent apoptotic phenotype. Transcripts or processes increased in the absence of Pacman are shown in blue text and those that reduce in *pacman* mutants are shown in red. Solid lines represent interactions that have been experimentally verified either by our group or by others. *Whilst the effect of Dilp8 on ecdysone has been shown, the hypothesised reduction in ecdysone in *pacman* mutants remains to be determined. Grey dashed lines represent unknown interactions including the factor (X) that is required for the apoptotic- and Pacman-mediated decrease *lncRNA:let7C* expression.

## DISCUSSION

In this study, we have used miRNA-seq in a global approach to identify candidate targets of the exoribonucleases Dis3L2 and Pacman during wing imaginal disc development. The use of well-characterised null mutations plus their isogenic controls has allowed us to assess the effects of these exoribonucleases on mature miRNA expression as well as reveal insights into the global expression of miRNAs in wing imaginal discs in a natural tissue. Our studies also allowed us to increase our understanding of the contribution of Pacman in inhibiting apoptosis in WIDs via the *let-7* miRNA complex. This work shows that cytoplasmic 5’-3’ RNA degradation plays a pivotal role in maintenance of the correct levels of miRNAs to regulate a suite of RNAs involved in this important cellular event.

Analysis of our miRNA-seq data revealed, for the first time, the global landscape of miRNA expression in relation to their genomic location. Interestingly, most dectected miRNAs are located within mRNAs (54%), although a substantial proportion are located within lncRNAs (27%). Of note, miRNAs encoded within lncRNAs, together with their host lncRNAs, tend to be more highly expressed than other miRNAs with their host lncRNAs also being longer and more abundant than average. The molecular mechanisms underlying these effects are not known. This data also shows that mirtrons are often less well expressed than other miRNAs, as previously observed (Berezikov et al. 2011).

The 3’-5’ exoribonuclease Dis3L2 was shown to have a minimal effect on the miRNA expression profile, however, we did validate an increase in the expression of *miR-4911-3p.* Interestingly, the *miR-4911* pre-miRNA is encoded within the second intron of the gene *ND23,* and therefore belongs in the category of miRtrons. Mirtrons have been shown in *Drosophila* to be preferentially uridylated by the TUTase Tailor and subsequently degraded by Dis3L2 (Bortolamiol-Becet et al. 2015; Reimao-Pinto et al. 2015). Tailor preferentially uridylates transcripts ending in a guanine (which is the 3’-most nucleotide following splicing), which is indeed the case for *miR-4911-3p* and the addition of uridyl residues to the G-terminal mirtrons produces a preferential substrate for Dis3L2, therefore our findings are entirely consistent with a role for Dis3L2 in degradation of this mirtron derived miRNA.

Previous work in *Drosophila* S2 cells revealed that the depletion of Tailor results in an increase of a substantial number of miRNAs (35%; 28 out of the 79 detected) (Reimao-Pinto et al. 2015). If uridylation by Tailor occurs at the same levels in wing imaginal discs, and the uridylated transcripts are predominantly degraded by Dis3L2, then we would expect to see upregulation of more miRNAs in *dis3L2* mutant imaginal discs. However, we see very few changes in miRNA expression which may suggest the wing imaginal disc is either less dependent upon this pathway for miRNA decay, or, more plausibly, there is redundancy between the miRNA decay pathways. For example, our previous work showed that loss of Dis3, another 3’-5’ exoribonuclease, which associates with the exosome, resulted in the altered expression of 39% of miRNAs in the wing imaginal disc (42 of 109 detected) (Towler et al. 2015). Therefore, although Dis3L2 affect the levels of pre-miRNAs, as has been shown in mouse and human cells, the levels of functional mature miRNAs appear to be less dependent upon Dis3L2 mediated decay. This is also consistent with other findings (Chang et al. 2013; Ustianenko et al. 2013; Nowak et al. 2017), although a more pronounced role was identified in immortalised human cells which also demonstrated the importance of 3’ terminal modification within the process (Shukla et al. 2019).

In contrast, loss of the 5’-3’ exoribonuclease Pacman has a much more profound effect on the levels of miRNAs in WIDs with 70 (32% of the total detected) being dysregulated, the majority of which (52) were upregulated. These results are consistent with those in *C. elegans* where depletion of xrn-1 resulted in accumulation of most of the miRNAs tested, including *let-7* (Chatterjee et al. 2011). In humans, XRN1 has been shown to regulate a subset of miRNAs in immortalised cell lines (HEK293T (Bail et al. 2010)) although the contribution of XRN1 in normal human tissue has not yet been tested. miRNAs transcribed from their own locus showed a trend of increased expression in the absence of Pacman. Interestingly, among the top 15 upregulated miRNAs, 5 (33%) were mirtrons suggesting that Pacman may be key in regulating the levels of these non-canonical miRNAs. Our studies also increase our understanding of the contribution of Pacman to apoptosis in wing imaginal discs and its subsequent effect on the *let-7* miRNA cluster. This work shows that cytoplasmic 5’-3’ RNA degradation plays a pivotal role in maintenance of the correct levels of miRNAs to regulate a suite of RNAs involved developmental timing and maintaining cellular homeostasis.

The biological function of the top 15 miRNAs is unknown, except for *miR-310*, *miR-311*, *miR-312* and *miR-313*, which are expressed as a cluster. Since both the 3p and 5p arms of *miR-312* are upregulated, the effect of Pacman on these miRNAs is presumably indirect and results from the transcriptional activation of the locus. All four members of this *miR-310* cluster have been experimentally shown to regulate the Toll-mediated immune response via inhibiting the expression of Drosomycin in flies infected by bacteria (Li et al. 2017).

The most significant finding in this paper is to illuminate the ways that Pacman regulates miRNAs to control apoptosis. The key finding is that Pacman controls the expression of the *lncRNA*:*let7C,* which encodes the conserved miRNAs *miR-100*, *let-7* and *miR-125* which in turn regulate development and apoptosis. Interestingly, *miR-277*, *263a* and *263b*, which target the 3’UTR of the pro-apoptotic RNA *hid* are also downregulated in *pcm* null mutants. Therefore, the data in this paper, together with our previous work, allows us to construct a model which may explain the effects of Pacman on apoptosis in wing imaginal discs (Fig 7) although further work is required to identify the intermediary factor(s) which govern transcriptional regulation, with Ecdysone being a promising candidate.

In summary, our experiments, using natural tissue rather than immortalised cell lines, show that the 5’-3’ exoribonuclease Pacman/XRN1 plays an important role in controlling the levels of mature miRNAs. Pacman also regulates the levels of mRNAs, lncRNAs and miRNAs (directly and indirectly) in the apoptosis pathway, linking the molecular effects on RNA levels with the Pacman phenotypes observed. Further studies should reveal the molecular mechanisms whereby this exoribonuclease specifically controls the levels of RNAs in wing imaginal discs to allow correct development and maintain homeostasis.

## MATERIALS AND METHODS

### Drosophila husbandry

*Drosophila* stocks were cultured on standard media and all experiments were performed at 25°C. *dis3L2^12^* and its isogenic control were produced in (Towler et al. 2020) and *pcm^14^* and its isogenic control were generated in (Waldron et al. 2015). The following stocks were obtained from Bloomington Stock Center: *nubbin-GAL4* (stock 25754; *P{UAS-Dcr-2*.*D}1*, *w^1118^; P{GawB}nub-AC-62*), *69B-GAL4* (stock 1774; *w*;; P{GawB}69B*), *H99* (stock 1576; *Df*(*3L*)*H99 kni^ri-1^ p^p^*/*TM3 Sb^1^), UAS-rpr* (stock 5824; *w^1118^; P{w[+mC]=UAS-rpr.C}14)*. *UAS-dilp8* was a kind gift from Dr Pierre Leopold.

### Wing imaginal disc dissection

Imaginal discs were dissected from 3rd instar wandering larvae; for *pcm^WT^*, *dis3L2^12^* and *dis3L2^WT^* 90 wing discs were dissected and pooled for each of the 4 replicates; whereas 180 discs were collected for each pool for the *pcm^14^* mutant, as they are 45% the size of controls (Waldron et al. 2015). Since *pcm* is on the X chromosome and the *pcm^14^* mutation is completely lethal at the pupal stage, the only mutant larvae available are hemizygous males. To account for any sex-specific differences in gene expression, male *pcm^WT^* larvae were collected. To allow accurate developmental staging and avoid overcrowding, 3 hour egg lays were performed. 120 hours after egg lay, wandering 3rd instar (L3) larvae were collected and dissected in Ringers solution (3 mM CaCl2, 182 mM KCl, 46 mM NaCl, 10 mM Tris pH7.2) under a Nikon SMZ800 dissection microscope and flash frozen in liquid nitrogen.

### miRNA-seq and bioinformatic analysis

RNA extraction was performed from between 60 and 180 120hr wing imaginal discs using the miRNeasy micro kit (Qiagen #217084) according to manufacturer’s instructions. *2S rRNA* depletion was performed essentially as described in (Aspden et al. 2014). Briefly, 60 µL of beads (Dynabeads MyOne Streptavidin C1, Thermo Fisher #65001) for each sample were bound to 750 pmoles of *2S rRNA* oligo in Tris-EDTA buffer. 30 µL of oligo-bound beads were used to deplete 400 ng of sample RNA. Two rounds of depletion were performed. Then, the RNA was precipitated and its integrity verified using an RNA 6000 nano chip (Agilent, #5067-1511) on a Bioanalyser 2100. sRNA-seq libraries were made using the QIAseq® miRNA library preparation kit (Qiagen #331502), according to the manufacturer’s instructions, using 6 µL of 2S-depleted RNA. Library quality was assessed using a DNA High Sensitivity chip (Agilent #5067-4626) on a Bioanalyser 2100. Libraries were sequenced on a NextSeq500 at Leeds Genomics obtaining a total of 326,514,877 reads with individual sample coverage ranging from 15.3 to 23.9 million reads.

To analyse the sRNA-seq data, miRDeep2 was used to build a genome with the dmel-all-chromosomes-6.29.fasta file from FlyBase (Consortium et al. 2018). Reads were mapped in miRDeep2 (Friedlander et al. 2012), with the parameters -h, -k, -l17 and -m and subsequently quantified again using miRDeep2. Differential expression was performed using DESeq2 using the default parameters in R. The data was independently analysed with sRNAbench from the sRNAtoolbox suite mapping (Aparicio-Puerta et al. 2019; Aparicio-Puerta et al. 2022) to BDGR6 (Hoskins et al. 2015) taking into account UMIs incorporated during the library preparation. To be classified as differentially expressed, miRNAs had to display a fold change >1.5 and padj<0.05 in the miRDeep2 analysis together with 4 of the 5 statistical outputs from sRNAbench. All normalised read count and fold change data presented in this work is from the miRDeep2 pipeline.

### RNA extraction and qRT-PCR

RNA extraction was performed from between 30 and 120 120hr wing imaginal discs or a single 120hr whole L3 larvae using the miRNeasy micro kit (Qiagen #217084) according to manufacturer’s instructions. Reverse transcription for miRNAs and *snoR442* were performed using the TaqMan miRNA reverse-transcription kit (Applied Biosystems #4366596) according to manufacturer’s instructions. Reverse transcription for other transcripts was performed using the hi-capacity cDNA reverse transcription kit (Applied Biosystems #4368814) according to manufacturer’s instructions. qRT-PCR-were performed in technical triplicate using the TaqMan Fast Universal PCR Master Mix (Thermo Fisher #4352042) according to manufacturer’s instructions in either a ViiA7 or QuantStudio7 Flex Real-Time PCR machine. All primers used in this study can be found in Supplemental File S2.

### Isloation of Ago-bound miRNAs

To isolate the miRNAs bound by Ago, the TraPR small RNA isolation kit (Lexogen #128) was used following manufacturers instructions. 60 *pcm^WT^* and 120 *pcm^14^* WIDs were used as starting material. The levels of the miRNAs were measured using TaqMan miRNA assays and normalised to the levels of *miR-275-5p* which showed consistent expression across all genotypes.

### Statistical Analysis

All statistical analyses were performed in either R v4.2.2 or GraphPad Prism 8. Two-sided two-sample t-tests were used to compare the means of single test groups to single control groups. If multiple comparisons were required, a one-way ANOVA was performed with a post-test to compare the means of each possible pair of samples. Outputs from statistical analyses can be found in Supplemental File 1 and Supplemental Tables S1-5.

## ACKNOWLEDGEMENTS

The authors wish to thank Dr Leandro Castellano, Dr Oliver Rogoyski and Dr Chris Jones for helpful discussions. We would also like to thank Clare Rizzo-Singh for technical help and Dr Pierre Leopold for providing the *UAS-dilp8* line. This work was funded by a BSMS studentship (WB003-46) to E.I.M.B and Biotechnology and Biological Sciences Research Council grants (BB/P021042/1 and BB/V001701/1) to S.F.N. and S.F.N plus B.P.T respectively.

## Author contributions

E.I.M.B designed and performed the experiments, analyzed much of the data and contributed to the writing of the manuscript. B.P.T. contributed to experimental design, performed experiments, analyzed and interpreted the RNA-seq data, prepared the figures, and contributed to the writing of the manuscript. S.F.N. coordinated the study, contributed to the design and interpretation of the experiments, and contributed to the writing of the manuscript.

## Supplemental Figure Legends

**Supplemental Fig S1: Descriptive statistics of miRNAseq in wing imaginal discs. A)** *dis3L2^WT^*and *pcm^WT^* miRNA expression show excellent correlations across all replicates. **B)** Correlation between the average normalised expression (CPM) of each detected miRNA between the two isogenic control lines demonstrates excellent correlation**. C)** miRNAs derived from host genes are globally more abundant than those transcribed from their own genomic locus. miRNA expression refers to the average CPM across all 8 isogenic control replicates. **D)** lncRNAs hosting miRNAs are longer than the genome average.

**Supplemental Fig S2: miRNA vs host relationships in ribonuclease mutants. A/B)** Boxplot displaying the fold change of miRNAs in *dis3L2^12^* **(A)** or *pcm^14^* **(B)** mutant wing discs by host RNA type. Dotted lines represent ^+^2 fold change. **C/D)** Proportion of miRNAs detected in wing discs and demonstrating ribonuclease sensitivity by genomic locus **(C)** or host RNA type **(D)**. A larger than expected proportion of miRNAs derived from their own locus show increased expression in ribonuclease mutants **. E/F)** Most changes in miRNA expression are independent of changes in their host RNA in either *dis3L2^12^* **(E)** or *pcm^14^* **(F)** mutant wing discs. Dotted lines represent ^+^2 fold change.

**Supplemental Fig S3: chinmo expression is increased in** pcm **mutant wing imaginal discs.** qRT-PCR validating increased *chinmo* expression in *pcm14* wing imaginal discs. n=3, error bars represent SEM, ***p<0.0002.

